# The Malate–Aspartate Shuttle supports thermogenic lipid mobilization in brown adipocytes

**DOI:** 10.1101/2025.08.04.667739

**Authors:** Michaela Veliova, Caroline M. Ferreira, Katrina Montales, Francisco Villalobos, Alexandra J. Brownstein, Rebeca Acín-Pérez, Gabrielly S. Ferreira, Linsey Stiles, Marc Liesa, Orian S. Shirihai, Marcus F. Oliveira

**Author notes:** These authors contributed equally to this work (Michaela Veliova, Caroline M. Ferreira). To whom correspondence should be addressed. Marcus F. Oliveira, Institute of Medical Biochemistry Leopoldo de Meis, CCS, Universidade Federal do Rio de Janeiro, A. Bauhinia 400, CEP 21941-590, Rio de Janeiro, RJ, Brazil. Orian S. Shirihai. Department of Molecular and Medical Pharmacology, Division of Endocrinology, Department of Medicine, David Geffen School of Medicine at UCLA, Los Angeles, CA, USA.

## Abstract

Brown adipose tissue (BAT) plays a central role in thermogenesis by coupling fatty acid oxidation to heat production. Efficient BAT thermogenic activity requires enhanced glycolytic flux, which in turn depends on continuous regeneration of cytosolic NAD⁺ to sustain glyceraldehyde-3-phosphate dehydrogenase activity. This regeneration is mediated by three main pathways: lactate dehydrogenase, the glycerol-3-phosphate shuttle, and the malate–aspartate shuttle (MASh). We previously showed that inhibition of the mitochondrial pyruvate carrier increases energy expenditure in brown adipocytes via MASh activation. However, the specific contribution of MASh to BAT energy metabolism remains poorly defined. Here, we show that MASh is functional and directly regulates lipid metabolism in BAT. Enzymatic activities of cytosolic and mitochondrial malate dehydrogenases and glutamic–oxaloacetic transaminases in BAT were comparable to those in the liver. Using a reconstituted system of isolated BAT mitochondria and cytosolic MASh enzymes, we demonstrated that extra-mitochondrial NADH is efficiently reoxidized in a glutamate-dependent manner via MASh. Genetic silencing of the mitochondrial carriers critical to MASh—namely the oxoglutarate carrier (OGC1) and aspartate–glutamate carrier (Aralar1) had no apparent effects on respiratory rates. However, silencing either OGC1 or Aralar1 led to the accumulation of small lipid droplets and impaired norepinephrine-induced lipolysis. Taken together, our data indicate a novel role of MASh in regulating BAT lipid homeostasis with potential implications to body energy expenditure and thermogenesis.

## Introduction

Brown adipose tissue (BAT) is a specialized thermogenic organ that dissipates energy as heat through a process known as non-shivering thermogenesis. This function is mediated by uncoupling protein 1 (UCP1), located in the inner mitochondrial membrane, which uncouples oxidative phosphorylation by dissipating the proton gradient. BAT activation in response to cold exposure or adrenergic stimulation triggers lipolysis through enzymes including adipose triglyceride lipase (ATGL) and hormone-sensitive lipase (HSL) [1,2] . This results in the release of free fatty acids (FFAs) from lipid droplets, which activate UCP. In addition, FFAs serve as substrates for β-oxidation and the tricarboxylic acid (TCA) cycle, thereby increasing nutrient oxidation and mitochondrial oxygen consumption [3]. Given BAT’s role in energy dissipation and glucose utilization, understanding its regulatory mechanisms is of growing interest in the context of obesity and metabolic disease [4].

Beyond lipid oxidation, thermogenically active BAT exhibits elevated glucose uptake and glycolytic flux [2]. One of the key glycolytic reactions—catalyzed by glyceraldehyde-3-phosphate dehydrogenase (GAPDH)—requires cytosolic NAD^+^ as a cofactor [5]. Therefore, a continuous supply of cytosolic NAD^+^ is essential for sustaining glycolysis and consequently, BAT thermogenesis [6]. Disruptions in this redox balance can compromise BAT function, highlighting the need to identify the metabolic systems that sustain NAD⁺ availability under thermogenic stress.

Since the mitochondrial inner membrane is impermeable to NADH [7], specialized pathways known as mitochondrial redox shuttles are required to transfer reducing equivalents from cytosolic NADH into the mitochondrial matrix [8,9]. By doing so, these shuttles simultaneously regenerate the cytosolic NAD^+^ essential for maintaining glycolytic flux. In mammalian cells, cytosolic NAD^+^ is regenerated by three primary mechanisms: lactate dehydrogenase (LDH), the glycerol-3-phosphate shuttle (GPSh), and the malate-aspartate shuttle (MASh). Interestingly, while LDH activity increases in the context f BAT activation, the GPSh appears to be inactive under adrenergic stimulation [10]. This leaves the MASh as a potentially crucial pathway for supporting cytosolic NAD^+^ regeneration during thermogenesis.

The MASh is a conserved cyclic redox pathway that circumvents the mitochondrial inner membrane’s impermeability to NADH by facilitating the transfer of reducing equivalents from the cytosol to the mitochondrial matrix through the coordinated action of two malate dehydrogenases (MDH), two glutamate-oxaloacetate transaminases (GOT), and two mitochondrial carriers (**Fig. 1A**). MASh operates by first using cytosolic NADH to reduce oxaloacetate to malate, a reaction catalyzed by MDH1. The resulting malate enters the mitochondrial matrix via the malate-alpha-ketoglutarate carrier (OGC1/SLC25A11). Inside the matrix, MDH2 oxidizes malate back to oxaloacetate, transferring the electrons to mitochondrial NAD^+^ to form NADH. To maintain the cycle, oxaloacetate and glutamate are converted to aspartate and α-ketoglutarate by GOT2. These products are then transported back into the cytosol via the aspartate-glutamate carrier (Aralar1/SLC25A12) and OGC1, where cytosolic glutamate-oxaloacetate transaminase 1 (GOT1) activity regenerates oxaloacetate from aspartate. By transferring electrons from glycolytic NADH to the electron transport system, the MASh enhances OXPHOS and contributes to maintaining a reduced NAD pool within the mitochondria [8,11–14]. Although MASh is a well-characterized metabolic pathway in many mammalian tissues [8,11], its specific function in BAT is not fully understood. This is particularly significant because BAT utilizes both lipids and glucose for thermogenesis, a process that demands strict control of the cellular redox state. Early studies demonstrated that MASh contributes to increased metabolic activity in BAT after birth, without being influenced by dietary protein intake [15]. Interestingly, cold exposure elevates the expression of key MASh components, including cytosolic and mitochondrial MDH and GOT1 [16]. More recent evidence demonstrates that induction of GOT1 by cold is a key mechanism for MASh activation in BAT, as this activation enhances fatty acid oxidation. Despite reduced glucose oxidation, increased glycolytic flux and glucose uptake support lipid utilization by providing oxaloacetate via pyruvate carboxylase (PC) during cold conditions [17,18].

**Fig. 1:**
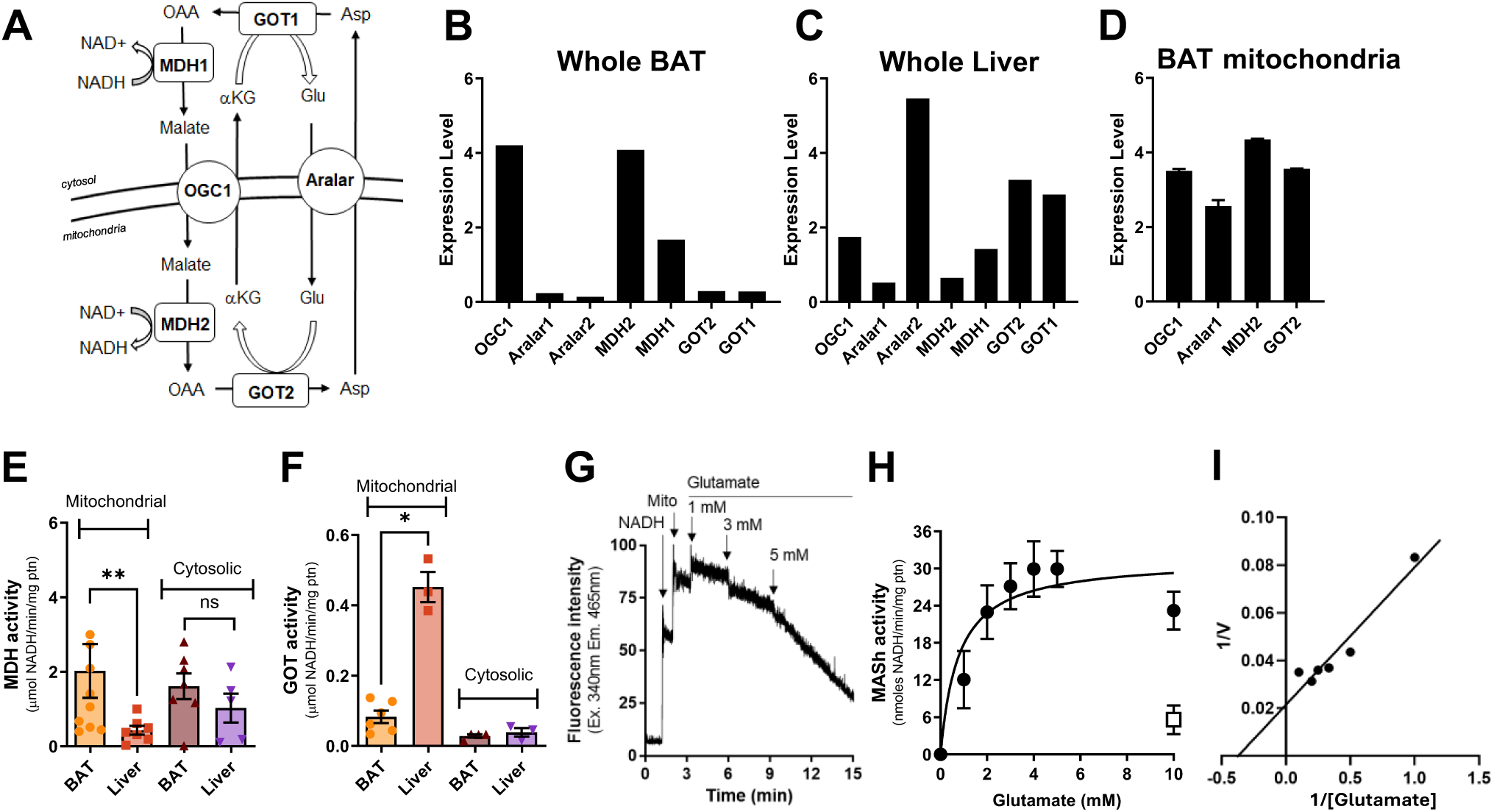
The Malate Aspartate Shuttle is active in brown adipocytes. (**A**) Schematic representation of the MASh. MDH, malate dehydrogenase, GOT, glutamic oxaloacetic transaminase; OGC1, oxoglutarate carrier 1; Aralar1, aspartate glutamate carrier 1; NAD^+^, nicotinamide adenine dinucleotide (oxidized form); NADH, nicotinamide adenine dinucleotide (reduced form). **(B-D**) *In silico* data mining analysis of the protein components of MASh whole BAT (**B**), and liver (**C**) determined in quantitative proteomic analyses conducted in [20]. (**D**) *In silico* data mining analysis of the protein components of MASh in BAT mitochondria determined in [21]. OGC: oxoglutarate carrier; Aralar: glutamate-aspartate exchanger; mMDH: mitochondrial malate dehydrogenase (MDH2); cMDH: cytosolic malate dehydrogenase (MDH1); mGOT: mitochondrial glutamate-oxaloacetate transaminase (GOT2); cGOT: cytosolic glutamate-oxaloacetate transaminase (GOT1). (**E**) Enzymatic activity measurements of cMDH and mMDH in BAT and liver. Data are expressed as mean ± SEM of at least five different experiments. (**F**) Enzymatic activity measurements of cGOT and mGOT in BAT and liver. Data are expressed as mean ± SEM of at least three different experiments. (**G**) Representative trace of NADH fluorescence during the assessment of MASh activity in BAT mitochondria. (**H**) MASh activity in BAT mitochondria in the presence of fluorocitrate (black circles) and fluorocitrate + aminooxyacetic acid (AOA) (white square). Data are expressed as mean ± SD of at least seven different experiments. (**I**) Lineweaver-Burke plot of data presented in (H).

Our previous work demonstrated that inhibiting the mitochondrial pyruvate carrier (MPC) in brown adipocytes increases mitochondrial respiration by activating the MASh [19]. This aligns with recent findings where MASh activation, via GOT1 overexpression, shifted cellular energy use from glucose to fatty acid oxidation [17]. Both observations suggest that MASh can compensate for metabolic restrictions by increasing the transfer of NADH into the mitochondria.

Based on this, we hypothesized that MASh plays a role in BAT metabolism beyond simply maintaining redox balance, potentially regulating lipid handling. In this study, we demonstrate that MASh is active in brown adipocytes and is crucial for lipid homeostasis by managing the balance between triglyceride storage and breakdown. While disrupting MASh at different points did not alter overall respiration in brown adipocytes, it triggered compensatory mitochondrial biogenesis and changed the dynamics of lipid droplets. These results establish MASh as a significant regulator of BAT metabolism and suggest that boosting its activity could be a promising strategy for improving metabolic health.

## Results

### The malate–aspartate shuttle is functional in murine brown adipose tissue

We previously demonstrated that pharmacological or genetic inhibition of the mitochondrial pyruvate carrier (MPC) in brown adipocytes enhances energy expenditure through the activation of lipid cycling and the malate–aspartate shuttle (MASh). Specifically, increased mitochondrial respiration under MPC inhibition was attenuated when expression of the MASh transporters, oxoglutarate carrier 1 (OGC1) and Aralar1, were downregulated—indicating that MASh is metabolically active in brown adipocytes [19]. Although earlier studies in rats also identified the activity of MASh enzymes in BAT [15], MASh’s broader significance and specific functions in BAT metabolism remain largely unexplored.

To characterize the presence and expression of MASh components in BAT, we first conducted an *in silico* analysis using a tissue-specific quantitative proteomic dataset from C57BL/6J mice [20,21]. We detected all six canonical components of the MASh in both brown adipocytes as well as in isolated mitochondria (**Fig. 1B, D**). Importantly, their expression levels were comparable to those found in the liver (**Fig. 1C**), a tissue where MASh function is well-established [22].

To determine if MASh is functional in BAT, we measured the enzymatic activities of MDH and GOT, in both cytosolic and mitochondrial fractions. Using liver tissue as a positive control, we found that mitochondrial MDH (MDH2) activity was approximately three-fold higher in BAT than in liver, whereas mitochondrial GOT (GOT2) activity was significantly lower (**Fig. 1E, F**). The cytosolic activities of both enzymes were comparable between the two tissues. These data support the existence of a functional MASh system in brown adipocytes.

To directly assess the functionality of MASh, we utilized a modified version of previously established methods [14,23,24]. This technique involves isolated BAT mitochondria reconstituted with the cytosolic components of the shuttle: MDH, GOT, NADH, aspartate, and glutamate. This experimental setup allows for the measurement of extra-mitochondrial NADH oxidation over time. The addition of glutamate initiates the cycling of metabolites, facilitating the transfer of reducing equivalents from the cytosol to the mitochondria via the MASh [24]. **Fig. 1G** shows that sequential additions of glutamate drive extramitochondrial NADH oxidation in a dose-dependent manner.

Glutamate titration revealed a classical Michaelis-Menten saturable curve, with an ^app^V_max_ of 31.5 nmol NADH/min/mg protein and an ^app^Km of 0.77 mM glutamate (**Fig. 1H, I**). These kinetic parameters were in the same range to those reported for mice brains in the absence of Ca^2+^ (^app^V_max_ = 38 nmol NADH/min/mg protein and ^app^K_m_ of 2.7 mM glutamate) [24,25]. Glutamate-induced NADH oxidation was strongly inhibited by the classical transaminase inhibitor aminooxyacetate (AOA), reinforcing the functionality of MASh in BAT (**Fig. 1H**). Collectively, these data directly demonstrates that MASh is active in BAT and unequivocally demonstrates its ability to transfer cytosolic reduced equivalents of NADH to brown adipocyte mitochondria.

### Genetic ablation of malate-aspartate-shuttle does not affect brown adipocyte energy expenditure

We next investigated the role of the MASh in brown adipocyte energy expenditure during thermogenic stimulation. To accomplish this, we silenced the two mitochondrial carriers essential for the MASh, Ogc1 and Aralar1, in primary brown adipocytes using adenovirus-mediated shRNA and siRNA, respectively (**Fig. 2A**). The knockdown efficiency for Aralar1 (Aralar1 KD) was approximately 80%, confirmed by qPCR. Importantly, this genetic intervention had no apparent effect on brown adipocyte differentiation, as the expression levels of six key differentiation markers were unaltered (**Fig. S1A**). Similarly, adenoviral-mediated knockdown of Ogc1 (Ogc1 KD) achieved an approximate 80% reduction compared to cells transduced with a scramble control. The expression of most differentiation markers, including *Ucp1, Elovl3,* and *Prdm16*, was unaffected. However, we observed significant increases in *Atgl, Pgc1α*, and *Tfam* mRNA levels, suggesting a partial reprogramming of the brown adipocyte differentiation pathway in Ogc1 KD cells (**Fig. S1B**).

**Figure 2:**
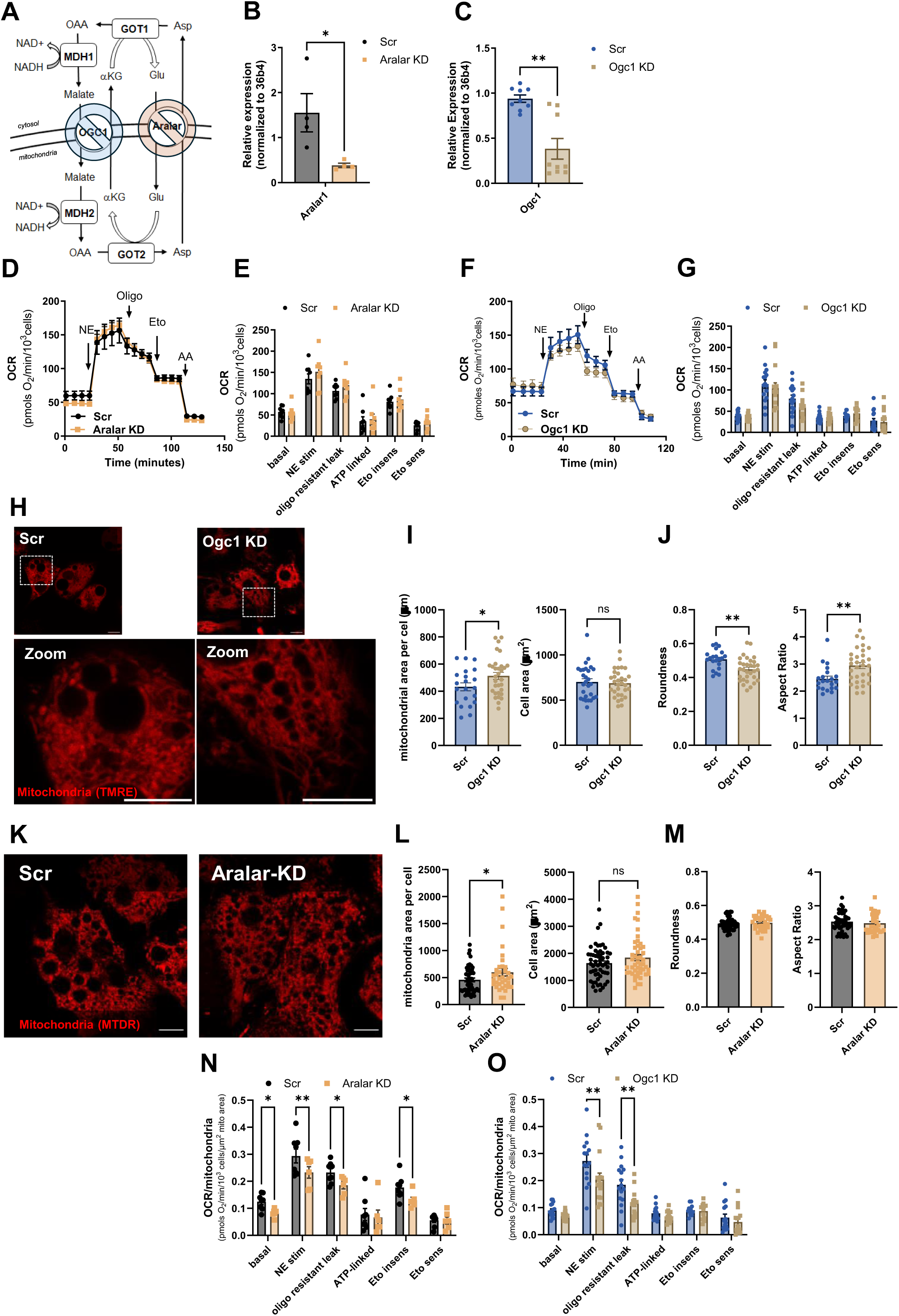
Genetic ablation of Malate-Aspartate-shuttle does not affect energy expenditure in activated brown adipocytes. (**A**) Schematic representation of the malate aspartate shuttle (MASh) and the two key mitochondrial transporters genetically silenced in this study. Adevirus-mediated shRNA was used to silence OGC1 (in blue), and siRNA transfection was used to silence Aralar1 (in orange). (**B**) mRNA levels of Aralar1 in brown adipocytes transfected with scramble RNA (Scramble) or Aralar1 siRNA (Aralar KD) from n=3 individual experiments. mRNA levels were normalized to 36b4. (**C**) mRNA levels of Ogc1 in brown adipocytes transduced with adenovirus containing scramble RNA (Scr) or shRNA for ogc1 (Ogc1 KD) from n=3 individual experiments. mRNA levels were normalized to 36B4. (**D and E**) Effect of Aralar1 silencing on cellular energy expenditure. Absolute oxygen consumption rates (OCR) were measured in respirometry media supplemented with 5 mM glucose and 3 mM glutamine. Norepinephrine (NE; 1 µM), oligomycin A (Oligo; 4 µM), etomoxir (Eto; 40 µM) and antimycin A (AA; 4 µM) were injected where indicated. (**D**) Representative OCR traces averaging 4 technical replicates (**E**) Quantification of non-stimulated, NE-stimulated, oligomycin-resistant leak, ATP-linked respiration, etomoxir sensitive and insensitive OCR from n=8 individual experiments. (**F and G**) Effect of Ogc1 silencing on cellular energy expenditure. Absolute oxygen consumption rates (OCR) were measured in respirometry media supplemented with 5 mM glucose and 3 mM glutamine. Norepinephrine (NE; 1 µM), oligomycin A (Oligo; 4 µM), etomoxir (Eto; 40 µM) and antimycin A (AA; 4 µM) were injected where indicated. (**F**) Representative OCR traces averaging 4 technical replicates. (**G**) Quantification of non-stimulated, NE-stimulated, oligomycin-resistant leak, ATP-linked respiration, etomoxir sensitive and insensitive OCAR from n=8 individual experiments. (**H**) Representative live-cell confocal images of primary brown adipocytes transduced with adenovirus containing either Scramble RNA (Scramble) or shRNA for ogc1 (OGC1 KD). Mitochondria were stained with 15 nM TMRE. Scale bar represents 10 µm. (**I**) Quantification of mitochondrial area per cell, and average cell area from images shown in (H). Data represent n=29-31 cells from 3 individual experiments. (**J**) Quantification of the effects of Ogc1 silencing on mitochondrial morphology parameters (roundness and aspect ratio). Data represent n=24-31 cells from 3 individual experiments. **(K)** Representative live-cell super-resolution confocal images of primary brown adipocytes transfected with either Scramble RNA (Scramble) or siRNA for Aralar1 (Aralar1 KD). Mitochondria were stained with 500nM mitotracker deep red (MTDR) prior to imaging. Scale bar represents 10 µm. (**L**) Quantification of mitochondrial area per cell, and average cell area from images shown in (K). Data represent n=45-55 cells from 3 individual experiments. (**M**) Quantification of the effects of Aralar1 silencing on mitochondrial morphology parameters (roundness and aspect ratio). Data represent n=54-55 cells from 3 individual experiments. **(N and O)** Quantification of the respiratory states determined in Aralar 1 (D-E) and Ogc1 (F-G) silenced brown adipocytes normalized by their respective mitochondrial content (I and L).

To determine whether Aralar1 KD or Ogc1 KD affects energy expenditure, we assessed respiration of intact brown adipocytes using Seahorse XF24 extracellular flux analysis. Oxygen Consumption Rate (OCR) measured by Seahorse provides a reliable proxy for cellular energy expenditure, as mitochondrial respiration drives most energy use in BAT. This method enables real-time assessment of mitochondrial function in intact cells with minimal stress. Measurements were performed in media containing 5 mM glucose and 3 mM glutamine and included basal respiration followed by norepinephrine (NE) stimulation to assess activated metabolic states. The glutamine concentration was chosen according to our previously established protocols [19], given its essential role as a precursor for glutamate, a key metabolite in the MASh pathway. Moreover, CPT1 inhibitor etomoxir was injected to assess the proportion of OCR that is dependent on fatty acid oxidation. Surprisingly, neither Aralar1 KD nor Ogc1 KD altered the mitochondrial metabolic parameters assessed (**Fig. 2 D-G**). These results underscore that the brown adipocyte differentiation program was not altered by the genetic interventions, since the level of norepinephrine-induced activation of oxygen consumption in both Aralar1 KD and Ogc1 KD were unchanged as previously indicated (**Fig. S1 A, B**). These findings indicate that MASh is dispensable for maintaining energy expenditure in activated brown adipocytes, possibly due to compensatory mechanisms.

Given that expression of *Tfam* and *Pgc1α*, markers of mitochondrial biogenesis, increased in Ogc1 KD brown adipocytes (**Fig. S1B**), we next examined whether mitochondrial content and morphology were affected. To accomplish this, we stained non-stimulated brown adipocytes with either TMRE (for Ogc1 KD cells) or mitotracker deep red (MTDR, for Aralar1 KD cells) and observed the fluorescence pattern by confocal microscopy (**Fig. 2H, K**) followed by digital image analyses. We observed that both Ocg1 KD cells as well as Aralar1 KD cells had increased mitochondrial area per cell compared with each of their scramble controls, without changing the total cell size (**Fig. 2I, L**). Interestingly we also found that mitochondria of Ogc1 KD cells had increased aspect ratio and reduced roundness, indicative of mitochondrial elongation (**Fig. 2J**). However, we did not observe differences in mitochondrial morphology in Aralar KD cells compared to their controls (**Fig. 2M**).

Together our results indicate that while the MASh is active in brown adipocytes, its disruption does not impair mitochondrial oxygen consumption. Instead, MASh-deficient cells seem to initiate a compensatory increase in mitochondrial biogenesis, likely to maintain thermogenic capacity by overcoming the reduced transfer of NADH equivalents. Indeed, knockdown of either Aralar1 or Ogc1 reduced basal oxygen consumption per organelle when normalized to mitochondrial content (**Fig. 2N, O**). This further supports the compensatory mitochondrial biogenesis triggered by disruption of the MASh. The altered mitochondrial morphology in Ogc1 knockdown cells (Fig. 2J) further suggests that these newly produced mitochondria are metabolically rewired, potentially due to changes in fuel availability [26].

### MASh disruption increases lipid droplet content and drives triglyceride storage in brown adipocytes

Given that Ogc1 KD upregulated *Atgl* and *Pgc1α* expression, we next investigated whether MASh impacts lipid metabolism. BODIPY 493/503 staining revealed that Ogc1 KD cells contained significantly more lipid droplets (LDs) than scramble transduced cells (**Fig. 3A, B**). Interestingly, LDs in Ogc1 KD cells were smaller in size compared to control (**Fig. 3C**). The total LD area, a surrogate of cellular lipid content, was not changed upon Ogc1 KD (**Fig 3D**). Importantly, when Aralar1 was silenced in brown adipocytes, we observed similar changes in LD number and size (**Fig. 3E-H**). This strongly indicates that MASh regulates LD dynamics in brown adipocytes.

**Figure 3:**
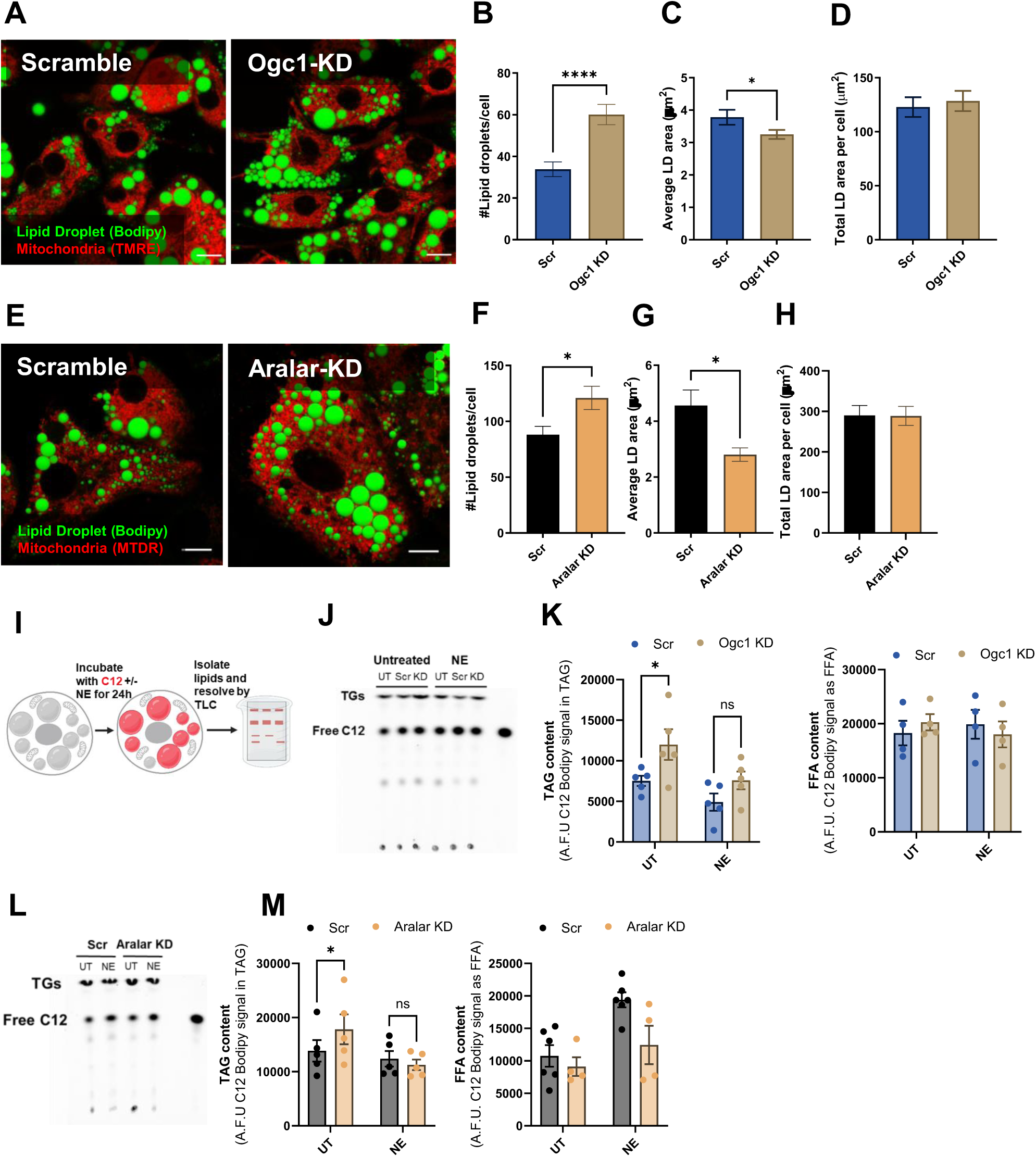
Knock-down of the Malate-Aspartate Shuttle alters Brown Adipocytes Triglyceride storage. **(A)** Representative live-cell confocal images of primary brown adipocytes transduced with adenovirus containing either Scramble RNA (Scramble) or shRNA for OGC1 (Ogc1 KD). Lipid droplets (LD) were stained with Bodipy 493 and mitochondria were stained with TMRE prior to imaging. Scale bar represents 10 µm. (**B-D**) Quantification of number of LD per cell, average LD area per cell, and total LD area per cell from multiple cells under the same conditions as images shown in (A). Data represent n=22-34 cells from 3 individual experiments. (**E**) Representative live-cell super-resolution confocal images of primary brown adipocytes transfected with either Scramble RNA (Scramble) or siRNA for Aralar1 (Aralar1 KD). Lipid droplets (LD) were stained with Bodipy 493 and mitochondria were stained with mitotracker deep red (MTDR) prior to imaging. Scale bar represents 10 µm. (**F-H**) Quantification of number of LD per cell, average LD area per cell, and total LD area per cell from multiple cells under the same conditions as images shown in (C). Data represent n=45-55 cells from 3 individual experiments. (**I**) Schematic representation of the experimental design for Figures J-M. Cells were incubated with Bodipy C12 (C12) for 24 hours in the presence and absence of 1 µM Norepinephrine (NE). Lipids were extracted from the cells and resolved by thin-layer chromatography (TLC). (**J**) Representative TLC plate of lipids extracted from primary brown adipocytes transduced with adenovirus containing either Scramble RNA (Scramble) or shRNA for ogc1 (OGC1-KD) in the presence and absence of NE. (**K**) Quantification of TAG from TLC samples shown in (J), n=4 individual experiments. (**H**) Representative TLC plate of lipids extracted from primary brown adipocytes transfected with either Scramble RNA (Scramble) or siRNA for Aralar1 (Aralar1 KD) in the presence and absence of NE. (**M**) Quantification of TAG from TLC samples shown in (H), n=4 individual experiments. Illustrations were created in BioRender.

Lipid droplet fragmentation can result from either increased triglyceride (TAG) synthesis or enhanced TAG breakdown. To test for increased esterification, we traced incorporation of a fluorescent fatty acid analog (BODIPY-C12) into lipids (**Fig. 3I**). After a 24 hr pulse and lipid extraction followed by thin-layer chromatography (TLC), both Aralar1 and Ogc1 KD cells showed elevated basal TAG content (**Fig. 3J-M**). Interestingly, under NE stimulation, TAG levels were similar across control, Aralar1 KD and Ogc1 KD. These results suggest that under non-stimulated conditions the MASh limits fatty acid esterification.

### MASh activity is essential for thermogenically activated lipolysis

We next developed a live-cell assay using fluorescently-labeled fatty acids to measure their mobilization from lipid droplets and assess the impact of MASh deficiency on adrenergically stimulated lipolysis, a key function of brown adipocytes. Brown adipocytes were pulsed with BODIPY-C12 for 12 hr, washed, and chased over 10 hr while neutral lipids were labeled with BODIPY 493/503. Imaging was performed using high-content confocal fluorescence microscopy (**Fig. 4A**). Control experiments validated the assay by showing that the BODIPY-C12 signal loss was accelerated by the lipolysis activators norepinephrine and forskolin, completely blocked by the ATGL inhibitor Atglistatin, and unaffected by the fatty acid oxidation inhibitor etomoxir, confirming the specific measurement of lipolysis (**Fig. 4B, C**). Applying this assay to Aralar1 KD cells completely abrogated norepinephrine-stimulated lipolysis compared to controls, consistent with a defect in acute lipolytic activation (**Fig. 4D, E**). To further validate this observation, we performed a longer pulse-chase assay with BODIPY-C12, followed by lipid extraction and TLC (**Fig. 4F**). Under NE treatment, control cells showed increased free fatty acid (FFA)-to-TAG ratios, indicative of lipolysis, whereas this response was lower in Aralar1 KD cells (**Fig. 4G, H**). These measurements truly reflect lipolysis as the presence of Atglistatin strongly reduces FFA-to-TAG ratios in both conditions (**Fig. 4H**). Finally, glycerol release into the media was used as a secondary measure of lipolysis. Norepinephrine treatment strongly increased glycerol release in control cells, but this response was partially blunted in Aralar1 KD adipocytes (**Fig. 4I**). Interestingly, glycerol release remained lower in Aralar1 KD cells than in controls even in the presence of Atglistatin, suggesting that MASh inhibition engaged alternative pathways of glycerol utilization (**Fig 4I**).

**Figure 4:**
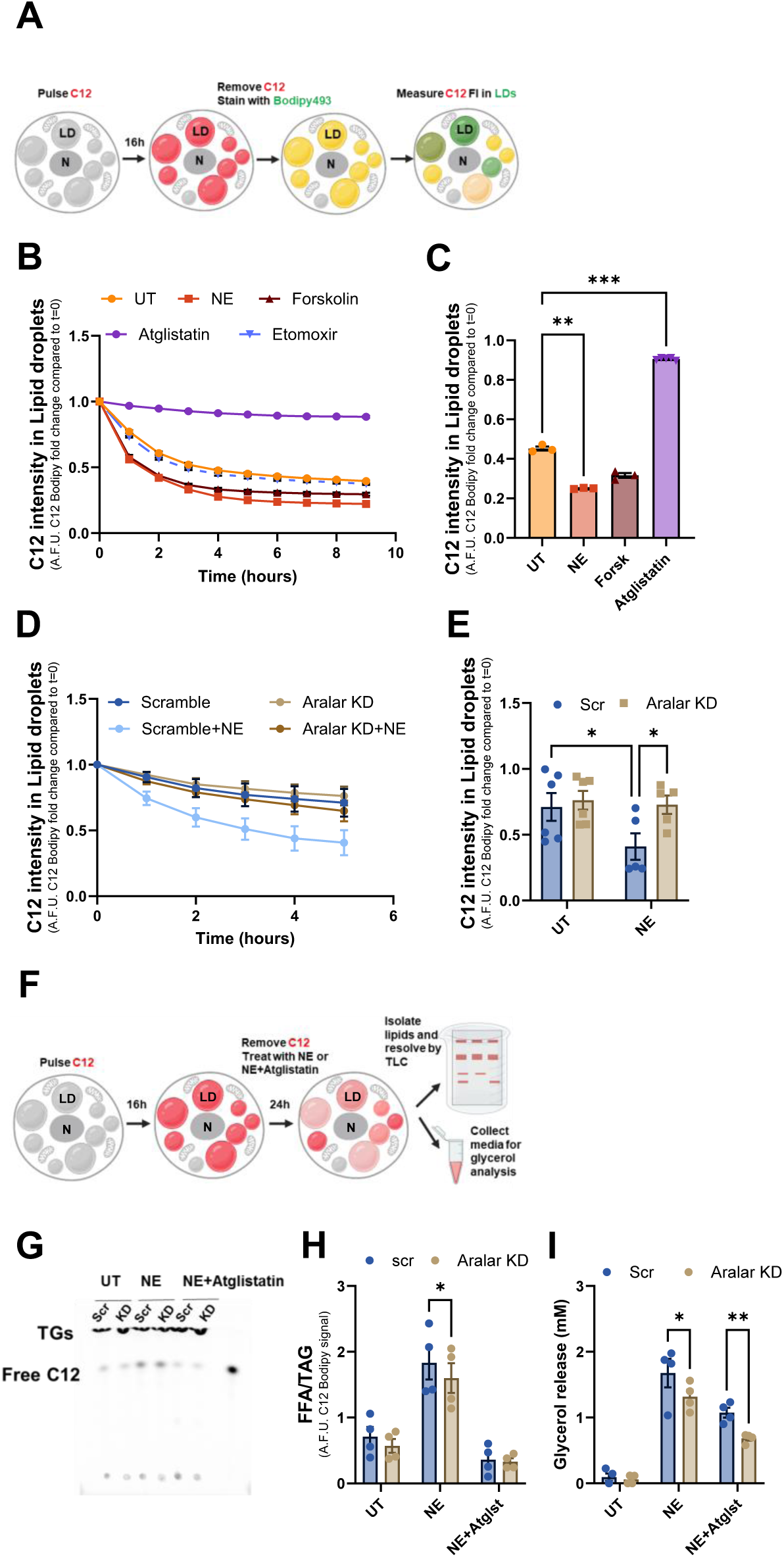
The Malate-Aspartate Shuttle plays a role in the lypolysis regulation of brown adipocytes. (**A**) Schematic representation of the experimental design for panels B-E. Primary brown adipocytes were incubated with Bodipy C12 (C12) for 16 hours to allow enough time for C12 esterification into TAGs. C12 was washed out and replaced by fresh media containing Bodipy 493 which stains all neutral lipids. Cells were imaged once per hour for a time-course of 5-9 hours on Operetta High-Content Imaging System. (**B**) Quantification of Bodipy C12 fluorescence intensity (FI) in lipid droplets, as determined by Bodipy 493 staining. Primary brown adipocytes were treated with either vehicle (UT), 1 µM Norepinephrine (NE), 10 µM Forskolin, 40 µM Atglistatin, or 10 µM Etomoxir. NE and Forskolin are known inducers of lipolysis in brown adipocytes, Atglistatin blocks lipolysis, and Etomoxir blocks fatty acid oxidation. Note that C12 signal decays faster in conditions where lipolysis is activated. Representative experiment averaging 4 technical replicates. (**C**) Quantification of C12 FI in lipid droplets after 5 hours of treatment. Data were normalized to t=0 and represent 3 individual experiments. (**D**) Quantification of Bodipy C12 fluorescence intensity (FI) in lipid droplets, as determined by Bodipy 493 staining. Primary brown adipocytes were transfected with either Scramble RNA (Scramble) or siRNA for Aralar1 (Aralar1 KD) and imaged in the presence and absence of 1 µM Norepinephrine (NE). Representative experiment averaging 3 technical replicates. (**E**) Data shows C12 FI in lipid droplets after 5 hours of treatment and was normalized to t=0. Data represents n=6 technical replicates from 2 individual experiments. (**F**) Schematic representation of the experimental design for panels G-I. Primary brown adipocytes were transfected with either Scramble RNA (Scramble) or siRNA for Aralar1 (Aralar1 KD). Cells were incubated with Bodipy C12 (C12) for 16 hours to allow enough time for C12 esterification into TAGs. C12 was washed out and replaced by fresh media containing either 1 µM Norepinephrine (NE), 40 µM Atglistatin, or a combination of both. After 24 h media was collected to measure cellular glycerol release, and cells were collected for lipid extraction and analysis by thin-layer chromatography (TLC). (**G**) Representative thin-layer chromatography (TLC) plate of lipids extracted from primary brown treated as described in (F). (**G**) Quantification of free C12/TAG from TLC samples shown in (E), as a measure or lipolysis. Data represents n=4 individual experiments. (**I**) Quantification of glycerol released into the media from n=4 individual experiments. Illustrations were created in BioRender.

## Discussion

The malate-aspartate shuttle (MASh) has been extensively characterized in tissues with high metabolic flux, such as the liver and heart [11,27], yet, its physiological relevance in thermogenically active BAT has remained largely unexplored. Here we provide direct evidence that MASh is functional in brown adipocytes and supports lipolysis, especially under adrenergic stimulation. Our findings reveal a previously unrecognized role for the MASh in regulating lipid metabolism in brown adipocytes, extending its classical function in maintaining cytosolic NAD^+^ redox balance in other tissues.

While direct evidence for MASh function in BAT is emerging, some previous studies highlight its biological importance [15–17]. Early research in rat BAT demonstrated that extramitochondrial NADH oxidation provided by MASh activity increased in response to a low-protein diet [15]. Another study found that cold exposure upregulated the expression of key components involved in recycling cytosolic NAD^+^, including MDH1/2 and GOT1 for MASh, glycerol 3-phosphate dehydrogenases (GPD1 and GPD2) for the glycerol phosphate shuttle (GPSh), and LDH [16]. This strongly suggests that to support high glycolytic rates of BAT during thermogenic stimuli, multiple pathways are engaged to provide cytosolic NAD^+^ posed by glyceraldehyde-3-phosphate dehydrogenase (G3PDH) reaction in glycolysis. In addition, a more recent study found that GOT1 is essential for sustaining fatty acid oxidation during cold exposure [17]. Although our data did not show altered fatty acid– driven respiration under acute norepinephrine treatment (**Fig. 2D-G**), this discrepancy likely arises from differences in experimental design—specifically, acute *in vitro* stimulation versus the prolonged *in vivo* cold adaptation in the recent study [17]. Importantly, our results indicate that MASh disruption impairs lipid metabolism upstream of β-oxidation by limiting the release of free fatty acids (**Fig. 3 and 4**). This suggests that MASh supports thermogenesis not by directly fueling fatty acid oxidation, but by enabling the initial lipolytic step that makes these fatty acids available.

Mechanistically, the connection between MASh and lipolysis may be mediated through the regulation of cytosolic NAD redox balance given that MASh activity regenerates NAD⁺ from NADH in the cytosol (**Fig. 1G-H**). In this regard, previous evidence demonstrates that stimulating NADH oxidation significantly enhances mitochondrial fatty acid oxidation by activating AMPK and carnitine palmitoyltransferase (CPT), while simultaneously suppressing acetyl CoA carboxylase (ACC) activity [28]. Our results demonstrate that in MASh-deficient cells, lipolysis is inhibited resulting in the accumulation of TAG and lipid droplets in brown adipocytes (**Fig. 3 and 4**). Conceivably, shifting the NAD redox balance towards the reduced state in brown adipocytes might impair the AMPK signaling cascade which would then stimulate ACC activity and inhibit β-oxidation. Alternatively, the oxidized state also supports this concept, as acetyl-CoA derived from fatty acids can acetylate and thereby activate components of the MASh, which could be the mechanism by which MPC inhibition increases MASh activity [19,29].

In addition, MASh disruption reduces lipid droplet size, which may impair ATGL-mediated hydrolysis of intracellular triglycerides, given that ATGL has been shown to preferentially act on larger lipid droplets [30]. Moreover, the accumulation of triglycerides may reflect passive re-esterification of fatty acids that were not mobilized due to impaired hydrolysis, rather than an active increase in de novo esterification. Notably, the unexpected result of MASh inhibition driving fatty acid esterification (**Fig. 3K-M**) parallel to enhanced *Atgl* expression (**Fig. S1B)** might be explained by the fact that mRNA levels may not reflect actual protein levels or activity.

Our previous work established that inhibiting mitochondrial pyruvate carrier (MPC) in brown adipocytes triggers a compensatory activation of the MASh, which in turn increases energy expenditure [19]. Intriguingly, MPC inhibition also stimulates lipid cycling and fatty acid utilization. Complementing these findings, genetic deletion of the MASh component GOT1 has been shown to produce the opposite effect: an upregulation of glucose uptake, glycolysis, and expression of the mitochondrial pyruvate carriers MPC1 and MPC2 [17]. Supporting these observations, an early study demonstrated that pharmacological inhibition of MASh increases the lactate/pyruvate ratio as a compensatory attempt to maintain cytosolic NAD^+^ which then limits the mitochondrial pyruvate metabolism [31]. This creates a compelling picture of a reciprocal feedback mechanism. Blocking the mitochondrial pyruvate metabolism activates MASh, while blocking the MASh enhances mitochondrial pyruvate metabolism. Therefore, MASh appears to function as a metabolic switch, mediating a preference for fatty acid oxidation when active and for glucose metabolism when its function is limited, as recently demonstrated [17].

The findings presented here suggest that disrupting the MASh activates the glycerol-3-phosphate shuttle (GPSh), a pathway normally suppressed in BAT during norepinephrine stimulation [10]. Interestingly, GPSh activation by G3PDH1 overexpression has been shown to increase lipid content in BAT [32]. This pathway can produce glycerol-3-phospate (G3P) from DHAP via GPD1, providing substrates for triglyceride synthesis even under redox imbalance. Increased availability of G3P, coupled with reduced FFA oxidation, could shift the metabolic balance toward lipid storage. The hypothesis of the GPSh acting as a compensatory pathway for NAD^+^ regeneration is supported by studies in GOT1-deficient BAT, where it is required to maintain glycolysis during thermogenic challenges [17]. This same compensatory mechanism would explain the reduced glycerol release we observed in Aralar1 knockdown brown adipocytes (**Fig. 4I**). We propose that by blocking the MASh, Aralar1 silencing engages the GPSh, which increases the cellular demand for its substrate, G3P. The cell meets this demand by phosphorylating the free glycerol pool, thereby limiting its release. It is important to note that while glycerol release is commonly used as a proxy for lipolysis, it may underestimate total lipid turnover. This measurement primarily captures the complete hydrolysis of triglycerides into free fatty acids (FFAs) and glycerol, without accounting for partial hydrolysis or re-esterification processes. Importantly, glycerol can originate not only from TAG breakdown in lipid droplets (LDs), but also from the hydrolysis of intermediates such as monoacylglycerols (MAGs), or from incomplete esterification during de novo lipogenesis [33]. Supporting this, we observed that Atglistatin—an inhibitor of ATGL that blocks the initial step of lipolysis—was unable to fully suppress norepinephrine-stimulated glycerol release (**Fig 4I**), despite effectively preventing TAG degradation as assessed by TLC and high content imaging (**Fig 4D-H**). This discrepancy suggests that a fraction of glycerol release may arise from ongoing lipid cycling rather than direct lipolysis. Recent work from our group and others has highlighted that brown adipocytes engage in active lipid cycling, wherein fatty acids are continuously hydrolyzed and re-esterified [19,34]. This cycle is metabolically demanding and depends on sustained cytosolic NAD⁺ availability to drive glycolysis and glycerol-3-phosphate (G3P) synthesis. The reduced glycerol release observed in MASh-deficient cells might thus reflect a broader disturbance in lipid turnover.

Another potential mechanism that might be engaged as a compensatory alternative for providing cytosolic NAD^+^ is the enzyme Aimf2, identified as a novel and BAT-specific external NADH dehydrogenase-like enzyme that mediates cytosolic NADH oxidation, glycolysis and thermogenesis in BAT [35]. However, its involvement in the context of MASh inhibition needs to be elucidated. Moreover, the existence of a redox compensatory mechanism in BAT highlights MASh as a critical node modulating both lipid storage and stimulated lipolysis.

## Conclusions

In brown adipocytes, functional MASh is essential for efficient thermogenic lipolysis. Disrupting the MASh triggers a compensatory increase in mitochondrial content, but these new organelles are metabolically reprogrammed, resulting in lower oxygen consumption per mitochondrion. Importantly, thermogenically activated lipolysis in brown adipocytes depends on a functional MASh. Therefore, our data position MASh as a central player in coordinating cytosolic redox state with lipid mobilization in BAT. By linking NAD⁺ recycling to lipolytic capacity, MASh supports the metabolic flexibility required for efficient thermogenesis, which depends on the coordinated regulation of glucose and fatty acid metabolism in brown adipocytes. Although the precise mechanisms connecting MASh activity and lipolysis remain to be elucidated, our data show that disrupting MASh impairs lipolysis, highlighting its critical role in BAT metabolism. These findings highlight a redox– lipid axis that could be targeted to enhance BAT function in metabolic disorders such as obesity and insulin resistance.

## Materials and Methods

### *In silico* protein level analysis

Protein levels of the MASh components were analyzed in murine intact brown adipocytes, liver [20], and BAT isolated mitochondria [21]. Expression values represent the ratio Heavy/Light in quantitative proteomics using SILAC mice.

### Animals

The investigation with experimental animals was approved by the Experimental Animal Ethics Committee of the Federal University of Rio de Janeiro under the protocol A09/17-050-16 and by the ARC/IACUC of the University of California, Los Angeles. 4 to 5 week old wild-type male C57Bl/6J mice (from either Cemib-UNICAMP, Brazil or Jackson Laboratory, Bar Harbor, ME, USA) were fed standard chow diet and kept under controlled conditions (19–22°C and a 14:10 h light-dark cycle) until euthanasia performed by asphyxiation in a CO2 atmosphere and subsequent cardiac puncture. All experiments were performed in accordance with the Guide for Care and Use of Laboratory Animals of the NIH (NIH Publications No. 8023, revised 1978).

### Mitochondrial isolation

Interscapular and subscapular brown adipose tissues were dissected and deposed in a small volume of ice-cold SHE buffer (sucrose 250 mM; HEPES 5 mM; EGTA 2 mM, pH 7.2). Pieces were then gently homogenized in a 15 mL Potter-Elvehjem tissue grinder in a Teflon pestle with 10 mL of either ice-cold SHE buffer for downstream enzymatic measurements or in SHE buffer + 2% fatty-acid-free bovine serum albumin for shuttle activity assessment. The homogenate was filtered through two layers of gauze and centrifuged at 8500 x g for 10 min at 4°C. Following centrifugation, the cytosol-containing supernatant was removed and reserved, and the mitochondrial-containing pellet was resuspended in SHE buffer. Mitochondrial isolation was performed as previously described [36] and protein concentration was determined using a BCA protein assay (Thermo Fisher Scientific, Roskilde, Denmark) or the Folin method [37].

### Mitochondrial enzyme activities

Glutamic oxaloacetic transaminase (GOT) activity was measured in cytosolic and mitochondrial fractions as described [38]. Malate dehydrogenase (MDH) activity was measured in cytosolic and mitochondrial fractions by adding the sample (100 µg of protein) to a quartz cuvette containing 89 mM Tris pH 7.8, 5 µM rotenone, and 160 µM NADH. Reaction was started by adding 5 mM oxaloacetate and NADH absorbance (340 nm) was measured by a Shimadzu UV-72550 spectrophotometer (Shimadzu Scientific Instruments, Japan).

### Assessment of MASh activity

Mitochondria were incubated with or without 2.5 mM fluorocitrate for 10 min at 4°C. 200 µg of mitochondrial protein was added to a quartz cuvette containing 1 mL of MSK buffer (75 mM mannitol; 25 mM sucrose; 5 mM K_2_PO_4_; 20 mM Tris-HCl; 0.5 mM EDTA; 100 mM KCl; 0.1% BSA; pH 7.4), 4 U/mL GOT, 6 U/mL MDH, 5 mM aspartate, 5 mM malate, 0.5 mM ADP, 200 nM ruthenium red, 8 µM CaCl_2_ and 0.25 mM fluorocitrate at 37°C and MASh activity was assessed by monitoring NADH fluorescence and calculating NADH oxidation rates, as previously described [14,23,25].

### Primary brown adipocyte culture

Primary brown adipocytes were isolated and cultured as described [39,40]. In brief, BAT tissue was dissected from interscapular, subscapular, and cervical regions of four male mice and digested in 1.5 mg/ml Collagenase Type II (Worthington, Lakewood, NJ). The tissue was then filtered through a 100 µm and 40 µm mesh and centrifuged. Cells were resuspended in brown adipocyte culture media (25 mM glucose DMEM supplemented with 10 % newborn calf serum (Sigma-Aldrich, St. Louis, MO), 4 mM glutamine, 10 mM HEPES, 0.1 mg/mL sodium ascorbate, 100 U/mL penicillin, 100 µ/mL streptomycin) and plated in a 6-well plate. Pre-adipocytes were incubated in a 37°C, 8 % CO_2_ for 72 hours and then lifted using STEMPro Accutase (Thermo Fisher Scientific, Roskilde, Denmark), counted, and re-plated in the final experimental vessel. 24 hours later differentiation was induced by switching the media to BAT differentiation media (growth media supplemented with 1 µM rosiglitazone maleate (Sigma-Aldrich, St. Louis, MO) and 4 nM human insulin (Humulin R, Eli Lilly, Indianapolis, IN). Cells were differentiated for 7 days and media was changed every other day.

### Gene silencing

#### Adenoviral transduction

On day 3 of differentiation, adipocytes were incubated with 1.5 µL/mL of adenoviral preparation (10^9^ particles/ml) for 24 h in BAT differentiation media containing 1 µg/mL polybrene (hexadimethrine bromide, Sigma-Aldrich, St. Louis, MO). Adenovirus was removed 24 hours later. Experiments were performed on day 7 of differentiation. Ad-m*SLC25A11* and Ad-mKate2 shControl were generated and purchased from Welgen (Worcester, MA).

#### siRNA transfection

Undifferentiated pre-adipocytes were transfected with scramble RNA or Aralar1 (*Slc25a12*) siRNA using Lipofectamine RNAiMAX transfection reagent (Thermo Fisher Scientific, Roskilde, Denmark) according to manufacturer’s protocol. In brief, culture media was removed from cells and cells were incubated with Opti-MEM media (Thermo Fisher Scientific, Roskilde, Denmark), Lipofectamine 3000 reagent and 100 nM scramble RNA or siRNA for 4 hours. Then DMEM with 1 % fetal bovine serum (Thermo Fisher Scientific, Roskilde, Denmark) was added to the cells and incubated overnight. The next day media was replaced with differentiation media. Experiments were performed on day 7 of differentiation. The following siRNAs were used: ON-TARGETplus Non-targeting Pool (D-001810-10-05) and ON-TARGETplus Mouse Slc25a12 siRNA (L-064268-01-0005) from Dharmacon (Lafayette, CO)

### Respirometry

Seahorse experiments were performed on day 7 of differentiation. Prior to respirometry measurements, culture media was replaced with respirometry media (Seahorse XF Base medium (Agilent Technologies, Santa Clara, CA) supplemented with 5 mM glucose and 3 mM glutamine when indicated, and incubated for 30-45 minutes at 37°C (without CO_2_). The ports of the Seahorse cartridge were loaded with the compounds to be injected during the assay (50 μL/port) and the cartridge was calibrated. Oxygen consumption rates were measured using the Seahorse XF24-3 extracellular flux analyzer (Agilent Technologies, Santa Clara, CA). The following compounds were used for injections during the assay: 1 µM norepinephrine (Levophed), 4 µM oligomycin A (Calbiochem, San Diego, CA), 40 µM etomoxir (Sigma-Aldrich, St. Louis, MO), 4 µM antimycin A (Sigma-Aldrich, St. Louis, MO). After the assay, cells were fixed using 4 % paraformaldehyde (Thermo Fisher Scientific, Roskilde, Denmark). To normalize the data for possible differences in cell number, nuclei were stained with 1 µg/mL Hoechst 33342 (Thermo Fisher Scientific, Roskilde, Denmark) and nuclei were counted using the Operetta High-Content Imaging System (PerkinElmer, Waltham, MA).

### Live cell imaging

#### Super resolution confocal microscopy

Super-resolution live cell imaging was performed on a Zeiss LSM880 using a 63x Plan-Apochromat oil-immersion lens and AiryScan super-resolution detector with a humidified 5 % CO_2_ chamber on a temperature-controlled stage (37°C). Cells were differentiated in glass-bottom confocal plates (Greiner Bio-One, Kremsmünster, Austria). Prior to imaging, cells were stained with 1 µM Bodipy 493/503 (Bodipy) (Thermo Fisher Scientific, Roskilde, Denmark) and 500 nM MitoTracker deep red (MTDR) (Thermo Fisher Scientific, Roskilde, Denmark) for 1 h, or 15 nM TMRE for 30 min. MTDR was washed out and cells were imaged in regular culture media in the presence of Bodipy. Cells stained with TMRE were measured in the presence of TMRE. Bodipy was excited with 488 nm laser, MTDR was excited with 633 nm laser, and TMRE was excited with a 552 nm laser. Image Analysis was performed in FIJI (ImageJ, NIH). Image contrast and brightness were not altered in any quantitative image analysis protocols. For representative images, brightness and contrast were equivalently modified in the different groups compared, to allow proper representative visualization of the effects revealed by unbiased quantitation. Detailed image analysis protocols are available upon request.

#### High-content imaging of lipolysis

Lipolysis imaging was performed on an Operetta High-Content Imaging System (PerkinElmer, Waltham, MA) using a 20x lens with a humidity controlled cell culture incubator (5 % CO_2_ and 37°C). Cells were differentiated in a black-walled, clear-bottom 96-well microplate. On the 6^th^ day of differentiation, cells were incubated with 1 µM Bodipy 558/568 C12 (Bodipy C12) (Thermo Fisher Scientific, Roskilde, Denmark) overnight. The next day, Bodipy C12 was washed out and cells were stained with Bodipy 488/503. Cells were incubated with the following compounds: 10 µM Forskolin (Sigma-Aldrich, St. Louis, MO); 10 µM etomoxir (Sigma-Aldrich, St. Louis, MO); 40 µM Atglistatin (Selleck Chemicals, Houston, TX); 1 µM norepinephrine (Levophed). Cells were imaged immediately after addition of compounds for 5-9 hours, taking one image per hour. Analysis was performed using the Operetta integrated software. Lipid Droplet area was defined based on Bodipy 493/503 signal, and Bodipy C12 fluoresces intensity changes over time were quantified.

### Thin layer chromatography of intracellular lipids

Cells were seeded and differentiated in 12-well plates. For measurements of fatty acid esterification, cells were incubated with 1 µM Bodipy 558/568 C12 and 1 µM norepinephrine (when indicated) for 24 h. For measurements of lipolysis, cells were incubated with 1 µM Bodipy 558/568 C12 overnight. The next day Bodipy C12 was washed out and cells were treated with 40 µM Atglistatin or 1 µM norepinephrine (as indicated) for 6 hours. Thin layer chromatography of intra-cellular lipids (Rambold *et al*, 2015) with minor modifications. Intra-cellular lipids and lipids from media were extracted in 500 µL chloroform. Chloroform was evaporated using the Genevac EZ-2 Plus Evaporating System (Genevac, Ipswich, United Kingdom). Lipids were then dissolved in 10 µL Chloroform and 1 µL was spotted on a TLC plate (Silica gel on TLC Al foils, Sigma-Aldrich, St. Louis, MO). Lipids were resolved based on polarity in a developer solution containing ethylacetate and cyclohexane in a 2:1 ratio. TLC plates were imaged on the ChemiDoc MP imaging system (Bio-Rad Laboratories, Hercules, CA). Band densitometry was quantified using FIJI (ImageJ, NIH).

### Total glycerol assessment in media

Experiments were performed with media from TLC experiments assessing lipolysis. Total glycerol released into the media was measured using Free Glycerol Reagent (#F6428, Sigma-Aldrich, St. Louis, MO) according to manufacturer’s instructions.

### Gene expression

Primary brown adipocytes transduced with the adenovirus (OGC1-KD) or transfected siRNA (Aralar1-KD) were collected on the 7^th^ day of differentiation. RNA extraction was performed using the Direct-zol RNA Miniprep Plus Kit® (Zymo Research, Irvine, CA) according to the manufacturer’s instructions. 1 μg of RNA was used for cDNA synthesis by the High-Capacity cDNA Reverse Transcription Kit® (Applied Biosystems, Foster City, CA). Then, qPCR was performed with 0.4 ng/μl cDNA and 240 nM of each primer. Gene expressions were normalized by an endogenous control (36b4).

### Statistical analyses

Data were presented as mean ± SEM for all conditions. Comparisons between groups were done by one-way ANOVA with Tukey’s or Holm Sidak’s test for pairwise multiple comparisons. When appropriate, two-way ANOVA with Tukey’s multiple comparisons test was employed. Pairwise comparisons were done by two-tailed Student’s t-test. Differences of P < 0.05 were considered to be significant. All graphs and statistical analyses were performed using GraphPad Prism 10 for Windows (GraphPad Software, San Diego, CA).

### Disclosure of artificial intelligence large language models use

The authors utilized the AI language models Gemini Pro 2.5 and Perplexity to enhance the English language in this manuscript. When revisions were needed, the models were instructed with the prompt: “Revise the following section in order to make it clearer and with an English-native speaker writing”. The authors retained full responsibility for the manuscript’s content, and all AI-assisted revisions were critically evaluated to confirm that the original scientific information was preserved with precision and correctness.

## Acknowledgements

We would like to thank Dr. Nathanael Miller for assistance with image analysis. Dr. Evan Taddeo, Dr. Karel Erion, Dr. Michael Shum, Dr. Anthony E. Jones, and Dr. Ajit S. Divakaruni for helpful discussions and advice. Illustrations created with <u>BioRender.com</u>. O.S.S. is funded by NIH-NIDDK 5-RO1DK099618-02. M.F.O. is funded by Conselho Nacional de Desenvolvimento Cientıfico e Tecnológico (grants #229526/2013-6, #404153/2016-0, #303044/2017-9 and #308629/2021-3) the Fundação Carlos Chagas Filho de Amparo à Pesquisa do Estado do Rio de Janeiro (FAPERJ) (grants #E-26/102.333/2013, #E-26/203.043/2016 and #E-26/210.409/2019) and by the Coordenação de Aperfeiçoamento de Pessoal de Nıvel Superior–Brasil (CAPES)–Finance Code 001 and CAPES-PrInt. M.L. is funded by PID2021-127278NB-I00, funded by MCIN/AEI/ 10.13039/501100011033 and by “ERDF A way of making Europe”, by the European Union. C.M.F. is funded by fellowships from Conselho Nacional de Desenvolvimento Científico e Tecnológico (141311/2016-9) and Capes (PDSE - 88881.187835/2018-01). Special thanks to Barbara Cannon and Jan Nedergaard for guiding our labs into the field of thermogenesis and brown adipocyte biology.

## Conflict of interest

The authors declare no conflict of interest.

## Author contributions

Conceptualization; CMF, MV, ML, MFO, OSS, Investigation; MV, CMF, FV, AB, GSF, KM, RAP, LS, MFO, Writing-Original Draft; MV, CMF, MFO. Writing-Review & Editing; MV, CMF, AB, RAP, MFO, OSS, ML.

## Data availability statement

The complete set of raw data will be made available, upon reasonable request, by the corresponding author.

## Abbreviations

ACC: Acetyl CoA Carboxylase
AMPK: AMP-Activated Protein Kinase
Aralar1: Aspartate-Glutamate Carrier 1 (SLC25A12)
BAT: Brown Adipose Tissue
Ca²⁺: Calcium ion
CPT: Carnitine Palmitoyltransferase
DHAP: Dihydroxyacetone Phosphate
FFA: Free Fatty Acid
G3P: Glycerol-3-Phosphate
GPSh: Glycerol-3-Phosphate Shuttle
GOT: Glutamate-Oxaloacetate Transaminase
GOT1: Cytosolic Glutamate-Oxaloacetate Transaminase 1
LD: Lipid Droplet
LDH: Lactate Dehydrogenase
MDH: Malate Dehydrogenase
MPC: Mitochondrial Pyruvate Carrier
MASh: Malate-Aspartate Shuttle
NAD⁺: Nicotinamide Adenine Dinucleotide
NADH: Nicotinamide Adenine Dinucleotide
NE: Norepinephrine
OGC1: Oxoglutarate Carrier 1 (SLC25A11)
Pgc1α: Peroxisome Proliferator-Activated Receptor Gamma Coactivator 1-alpha
Prdm16: PR Domain Containing 16
TAG: Triacylglycerol (Triglyceride)
Tfam: Mitochondrial Transcription Factor A
TMRE;: Tetramethylrhodamine, Ethyl Ester
TLC: Thin Layer Chromatography
UCP1: Uncoupling Protein 1
GAPDH: Glyceraldehyde-3-Phosphate Dehydrogenase

## Notes

### Competing Interest Statement

The authors have declared no competing interest.

