## Supplementary table and figure for "The Malate–Aspartate Shuttle supports thermogenic lipid mobilization in brown adipocytes"

**Supplementary table 1.**

Primers used for quantitative PCR

| Gene | Forward primer 5'-3' | Reverse Primer 5'-3' |
|---|---|---|
| SCL25A11 | AGT CTC CTC TTG GGT GTT AGA | CTT CTG CTT TCT CCT GTC TCC |
| SCL25A12 | TGGTTACCTACGAGCTTCTGC | ACCGATGTGATCGGGGTTG |
| Ucp1 | GGC CTC TAC GAC TCA GTC CA | TAA GCC GGC TGA GAT CTT GT |
| Atgl | TCC GAG AGA TGT GCA AAC AG | CTC CAG CGG CAG AGT ATA GG |
| pgc1a | AAG ATC AAG GTC CCC AGG CAG TAG | TGT CCG CGT TGT GTC AGG TC |
| tfam | CAC CCA GAT GCA AAA CTT TCA | CTG CTC TTT ATA CTT GCT CAC AG |
| elov13 | TCC GCG TTC TCA TGT AGG TCT | GGA CCT GAT GCA ACC CTA TGA |
| prdm16 | GCC ATG TGT CAG ATC AAC GA | CCT TCT TTC ACA TGC ACC AA |
| 36b4 | GTC ACT GTG CCA GCT CAG AA | TCA ATG GTG CCT CTG GAG AT |

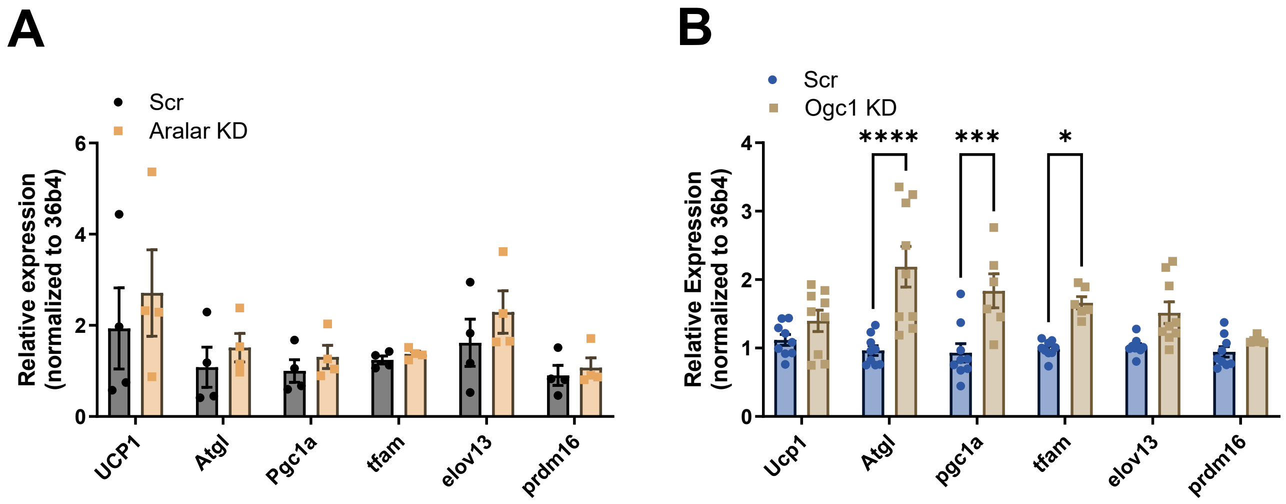


**Figure S1:** (**A**) mRNA levels of *Ucp1*, *Atgl*, Pgc1α, *Tfam*, *Elov13* and *Prdm16* in brown adipocytes transfected with scramble RNA (Scramble) or Aralar1 siRNA (Aralar KD) from n=3 individual experiments. mRNA levels were normalized to 36B4. (**B**) mRNA levels of *Ucp1*, *Atgl*, *Pgc1α*, *Tfam*, *Elov13* and *Prdm16* in brown adipocytes transduced with adenovirus containing scramble RNA (Scr) or shRNA for Ogc1 (Ogc1 KD) from n=3 individual experiments. mRNA levels were normalized to 36b4.
